# Assessing the effect of hypoxia on cardiac metabolism using hyperpolarized ^13^C magnetic resonance spectroscopy

**DOI:** 10.1101/495069

**Authors:** Lydia M. Le Page, Oliver J. Rider, Andrew J. Lewis, Victoria Noden, Matthew Kerr, Lucia Giles, Lucy J. A. Ambrose, Vicky Ball, Latt Mansor, Lisa C. Heather, Damian J. Tyler

## Abstract

Hypoxia plays a role in many diseases and can have a wide range of effects on cardiac metabolism depending on the extent of the hypoxic insult. Non-invasive imaging methods could shed valuable light on the metabolic effects of hypoxia on the heart *in vivo*. Hyperpolarized carbon-13 magnetic resonance spectroscopy (HP ^13^C MRS) in particular is an exciting technique for imaging metabolism that could provide such information.

The aim of our work was, therefore, to establish whether hyperpolarized ^13^C MRS can be used to assess the *in vivo* response of cardiac metabolism to systemic acute and chronic hypoxic exposure. Groups of healthy male Wistar rats were exposed to either acute (30 minutes), one week or three weeks of hypoxia. *In vivo* MRS of hyperpolarized [1-^13^C]pyruvate was carried out along with assessments of physiological parameters and ejection fraction. No significant changes in heart rate, respiration rate, or ejection fraction were observed at any timepoint. Haematocrit was elevated after one week and three weeks of hypoxia.

Thirty minutes of hypoxia resulted in a significant reduction in pyruvate dehydrogenase (PDH) flux, whereas one or three weeks of hypoxia resulted in a PDH flux that was not different to normoxic animals. Conversion of hyperpolarized [1-^13^C]pyruvate into [1-^13^C]lactate was elevated following acute hypoxia, suggestive of enhanced anaerobic glycolysis. Elevated HP pyruvate to lactate conversion was also seen at the one-week timepoint, in concert with an increase in lactate dehydrogenase (LDH) expression. Following three weeks of hypoxic exposure, cardiac metabolism was comparable to that observed in normoxia.

We have successfully visualized of the effects of systemic hypoxia on cardiac metabolism using hyperpolarized ^13^C MRS, with differences observed following 30 minutes and 1 week of hypoxia. This demonstrates the potential of *in vivo* hyperpolarized ^13^C MRS data for assessing the cardiometabolic effects of hypoxia in disease.

## Introduction

Oxygenation of tissue is key to survival and maintenance of organ health. The heart has the potential to be exposed to a spectrum of hypoxic insults, ranging from mild and transient, to prolonged and severe. The metabolic effects of acute hypoxia are well documented, and notably involve increased glycolytic flux and transient lactate acidosis^1,2^. Prolonged and severe hypoxia requires reprogramming of cardiac metabolism; the heart downregulates oxygen-consuming processes and upregulates glycolysis in an attempt to maximize ATP production under oxygen restricted conditions^3–5^. The effects of chronic hypoxia are observed in response to high altitude^6^, or as a factor in many pathological conditions; examples include chronic obstructive pulmonary disease^7^, complications in pregnancy^8^, sleep apnoea^9^, myocardial infarction (the peri-infarct region)^10^ and heart failure^11^.

However, much of this existing literature relies on *ex vivo* assessment of the metabolic changes that occur. As such, non-invasive *in vivo* measures of the effect of oxygen levels on cardiac tissue would be valuable, especially as the hypoxic response can be very transient^12^. Imaging techniques have begun to probe *in vivo* oxygen levels, and current prominent methods include blood-oxygen-level dependent (BOLD) MRI and positron emission tomography (PET) imaging, although neither is standard clinical practice as yet. BOLD MRI enables assessment of vascular oxygenation, using the paramagnetic nature of deoxyhaemoglobin to create image contrast^13^; this technique has not yet reached the clinic due to a combination of many challenges including low signal-to-noise and a need for robust analysis^14^, which studies have begun to address^15^. There is also an ongoing search for PET probes to assess hypoxia, the most promising of which is currently ^11^C-acetate. Clearance of this tracer is dependent on oxidative metabolism, and so accumulation indicates low oxygen presence^16^. It does, however, have a short half-life^17^ and so usage depends on a nearby cyclotron, and as with all PET options, patients will be exposed to ionizing radiation which may prohibit repeated measurements.

Spectroscopic imaging holds potential for providing non-invasive, non-radioactive metabolic data. Imaging of carbon-13 (^13^C) in particular can be very informative given the abundance of carbon present in metabolites, including those in pathways affected by oxygen level. Although ^13^C spectroscopy suffers from inherently low sensitivity *in vivo*, the advent of hyperpolarized ^13^C magnetic resonance spectroscopy (HP ^13^C MRS) offers the unique ability to measure the rate of enzyme flux *in vivo*^18^. It provides an enhancement of the ^13^C signal of >10,000 fold, and as such, enables a non-invasive measurement of enzymatic flux in real time. In the heart, the glycolytic pathway is central to the metabolic changes that occur as oxygen levels fall. The most established hyperpolarized ^13^C-labelled probe, [1-^13^C] pyruvate, is relevant to this pathway, as it allows us to visualise the fate of pyruvate either through mitochondrial pyruvate dehydrogenase (PDH) to bicarbonate, or through cytosolic lactate dehydrogenase (LDH) into lactate^19^. A previous study by Laustsen *et al.*^20^ showed the value of hyperpolarized pyruvate in the investigation of hypoxia in the diabetic rat kidney – demonstrating an ability to measure increased lactate production after fifteen minutes of hypoxic anaesthesia. Hypoxia is also one of many pathological factors of tumor development^21^, fluctuating over time and in regions of the tumor^22^, and as such Iveson *et al.* used HP ^13^C MRS in a mouse tumor model, showing that inspiration of a hypoxic atmosphere caused increased lactate production in tumors^23^. Oxidative stress has been investigated in a few non-cardiac studies, using HP dehydroascorbate^24,25^, but the toxicity of this compound may limit translation to clinical studies^26^. Indeed, the challenges and future of hyperpolarized probes for assessing renal and cardiac oxygen metabolism have been discussed in a review by Schroeder and Laustsen^27^. Thus far, no studies have investigated the use of HP ^13^C MRS to assess the effect of hypoxia on glucose metabolism in the *in vivo* heart.

In this study we have therefore assessed the effect of three lengths of hypoxic exposure – thirty minutes, one week, and three weeks - on the *in vivo* rat heart, using hyperpolarized [1-^13^C] pyruvate. We have measured the conversion of HP pyruvate to bicarbonate, lactate and alanine (**Figure 1A** shows the biochemical pathways involved). The level of oxygen saturation in the blood was matched across groups, and established following measurement in animals housed at 11% oxygen from previous rodent studies in our laboratory^3,4^. Alongside cardiac metabolism by MRS, we assessed ejection fraction by CINE MRI imaging, and measured heart rate and respiration rate in all groups. We further measured body weight and haematocrit in the longer exposure groups (1 week and 3 weeks hypoxia). In these latter groups, expression levels of cardiac PDH regulators pyruvate dehydrogenase kinase (PDK) 1, 2 and 4, and the expression level of lactate dehydrogenase (LDH), responsible for conversion of pyruvate to lactate, were also measured in cardiac tissue.

**Figure 1:**
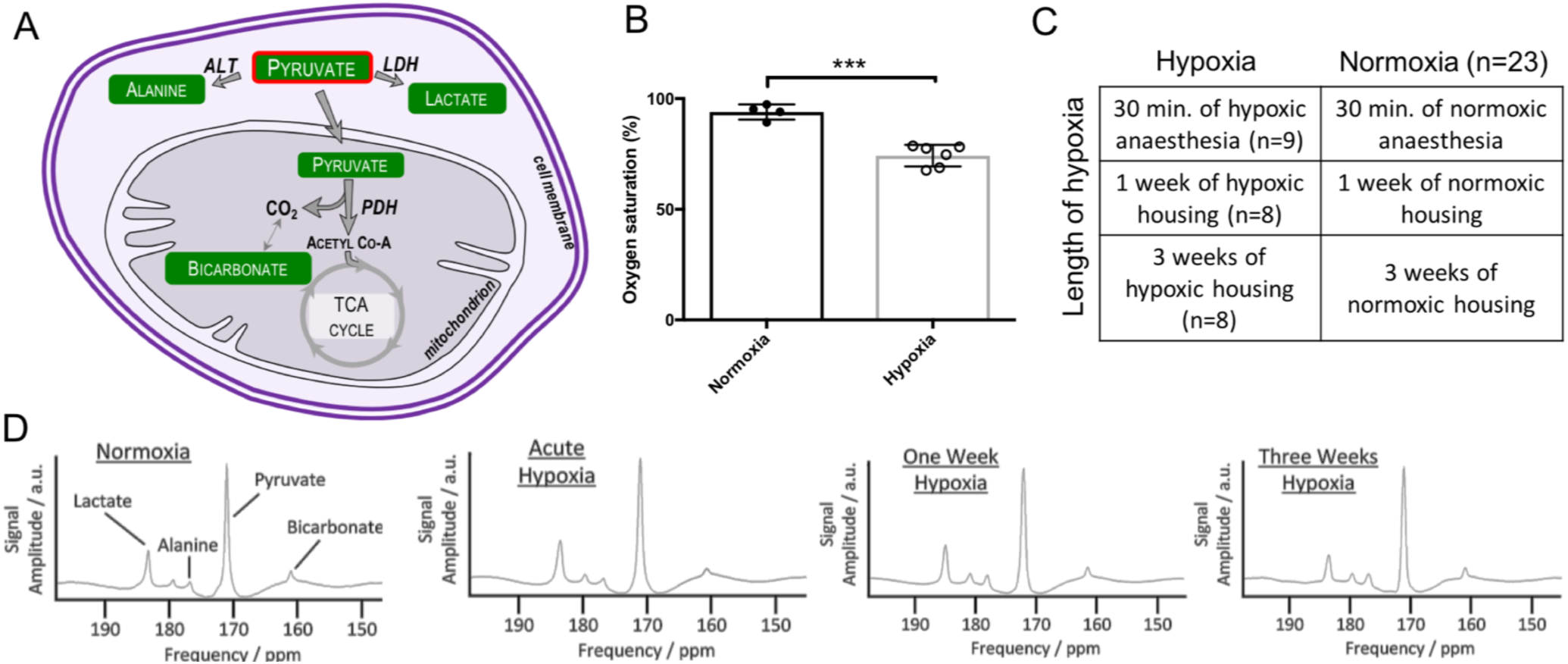
**(A)** Biochemical pathways visualized using HP [1-13C] pyruvate (outlined in red). **(B)** oxygen saturation of animals housed in the hypoxic chamber ***p=0.0001. **(C)** Experimental animal groups for three lengths of hypoxic exposure. Normoxic data subsequently treated as one group, n=23. **(D)** example summed spectra from each timepoint.

## Methods

### Animal handling

Male Wistar rats (initial body weight ∼200 g, Harlan, UK) were housed on a 12:12-h light/dark cycle in animal facilities at the University of Oxford. All imaging studies were performed between 6 am and 1 pm with animals in the fed state. All procedures conformed to the Home Office Guidance on the Operation of the Animals (Scientific Procedures) Act of 1986 and to University of Oxford institutional guidelines.

### Hypoxic exposure

A group of hypoxically-housed animals (n=6) and a group of animals housed in normoxia (n=4) were used to assess blood oxygen saturation. Saturation was measured to be 74±2% (**Figure 1B**) in hypoxia, using a pulse oximeter on their hind paw (MouseOx, Starr Life Sciences). This concentration was subsequently matched for all hypoxic exposures.

### Experimental groups for in vivo imaging

Three groups of animals were exposed to three different lengths of hypoxia. Control groups experienced normoxia only. The groups are summarized in **Figure 1C**.

### Thirty minutes (acute) hypoxia

Animals (n=9) were anaesthetised using isoflurane (2%) in 100% O_2_ (2L/min). Metabolic and functional data were acquired in normoxia as described in the imaging protocol below. Animals were then slowly introduced to hypoxia by increasing replacement of oxygen with nitrogen over thirty minutes, until a blood oxygen saturation which matched that of the animals housed in the hypoxic chamber was achieved (described above). A second injection of hyperpolarized [1-^13^C]pyruvate was administered and a second data set acquired. Acute hypoxia elicited some rapid physiological responses such as increased ventilation and heart rate^28^, which settled prior to data acquisition, allowing acquisition of data in a stable hypoxic state.

### One week of hypoxia

Animals (n=8) were housed in a normobaric hypoxic chamber for one week, during which time the oxygen concentration was reduced daily by 1-2% until at the final day the concentration was 11%. Animals were weighed daily, which resulted in brief exposure to normoxia (no longer than 5 minutes). Animals were subsequently anaesthetised under hypoxia (O_2_/N_2_ mix) outside the chamber, before being placed in the magnet and the imaging protocol. executed. A control group (n=6) was housed outside the hypoxic chamber in room air (21% oxygen) for one week from which normoxic data were acquired.

### Three weeks of hypoxia

Animals (n=8) were introduced to the normobaric hypoxic chamber as for the one-week experiments, but remained in the chamber for a further 14 days at 11% oxygen. Animals were then anaesthetised under hypoxia outside the chamber (O_2_/N_2_ mix) and underwent the MR protocol as for the one-week animals, to obtain *in vivo* cardiac metabolic data. A control group (n=8) was housed outside the hypoxic chamber in room air (21% oxygen) for three weeks from which normoxic data were acquired.

### Magnetic resonance (MR) protocol

Animals were anaesthetised with isoflurane (3.5% induction and 2% maintenance). Rats were positioned in a 7 T horizontal bore MR scanner interfaced to a Direct Drive console (Varian Medical Systems, Yarnton, UK), and a home-built ^1^H/^13^C butterfly coil (loop diameter, 2 cm) was placed over the chest. Correct positioning was confirmed by the acquisition of an axial proton fast low-angle shot (FLASH) image (TE/TR, 1.17/2.33 ms; matrix size, 64 x 64; FOV, 60 x 60 mm; slice thickness, 2.5 mm; excitation flip angle, 15°). An ECG-gated axial CINE image was obtained (slice thickness:1.6 mm, matrix size:128×128, TE/TR:1.67/4.6 ms, flip angle:15°) at the level of the papillary muscles for ejection fraction calculation. An ECG-gated shim was used to reduce the proton linewidth to ∼120 Hz. Hyperpolarized [1-^13^C]pyruvate (Sigma-Aldrich, Gillingham, UK) was prepared by 40 minutes of hyperpolarization at ∼1K as described by Ardenkjaer-Larsen *et al.*^18^, before being rapidly dissolved in a pressurised and heated alkaline solution. This produced a solution of 80 mM hyperpolarized sodium [1-^13^C]pyruvate at physiological temperature and pH, with a polarization of ∼30%. One millilitre of this solution was injected over ten seconds via a tail vein cannula (dose of ∼0.32 mmol/kg). Sixty individual ECG-gated ^13^C MR slice selective, pulse-acquire cardiac spectra were acquired over 60 s after injection (TR, 1 s; excitation flip angle, 5°; slice thickness 10 mm, sweep width 13,593 Hz; acquired points 2,048; frequency centred on the C1 pyruvate resonance)^29^.

### Tissue collection

All animals were sacrificed with an overdose of isoflurane following completion of the MR protocol. The heart was rapidly removed, washed briefly in phosphate buffered saline, and snap-frozen in liquid nitrogen.

### Blood analyses

Samples of blood were collected from the chest cavity on sacrificing, and centrifuged at 8,000 rpm for 10 minutes. Haematocrit was measured using a microhaematocrit reader (Hawksley, UK).

### Tissue analysis

For Western blotting of cardiac tissue from one week and three week groups, frozen tissue was crushed and lysis buffer added before tissue was homogenised; a protein assay established the protein concentration of each lysate. The same concentration of protein from each sample was loaded on to 12.5% SDS-PAGE gels and separated by electrophoresis^30^. Primary antibodies for PDK 1 and 2 were purchased from New England Biolabs and Abgent, respectively; an antibody for PDK4 was kindly donated by Prof. Mary Sugden (Queen Mary’s, University of London, UK). A primary antibody for LDH was purchased from Abcam (ab52488). Even protein loading and transfer were confirmed by Ponceau staining (0.1% w/v in 5% v/v acetic acid, Sigma-Aldrich), and internal standards were used to ensure homogeneity between samples and gels. Bands were quantified using UN-SCAN-IT gel software (Silk Scientific, USA) and all samples were run in duplicate on separate gels to confirm results.

### Magnetic resonance data analysis

All cardiac ^13^C spectra were analysed using the AMARES algorithm in the jMRUI software package^31^. **Figure 1D** shows example spectra summed over 30 seconds of acquisition in normoxic animals, acutely hypoxic animals and animals housed in hypoxia for one and three weeks, showing cardiometabolic conversion of the injected hyperpolarized pyruvate into the downstream products lactate, alanine and bicarbonate. Spectra were DC offset-corrected based on the last half of acquired points. The peak areas of [1-^13^C]pyruvate, [1-^13^C]lactate, [1-^13^C]alanine and [^13^C]bicarbonate at each time point were quantified and used as input data for a kinetic model based on that developed by Zierhut *et al.*^32^ and Atherton *et al*.^33^. PDH flux was quantified as the rate of ^13^C label transfer from pyruvate to bicarbonate. The rate of ^13^C label transfer from pyruvate to lactate and alanine was used as a marker of lactate dehydrogenase activity and alanine aminotransferase activity respectively. CINE images were analyzed using cmr42 software (Circle Cardiovascular Imaging, Calgary, Canada) by an experienced analyst blinded to experimental group.

### Statistical analyses

No significant differences were observed between the three normoxic groups (acute, one week and three weeks) for any parameter, therefore all normoxic values were combined for subsequent analysis. Values are reported as the mean ± standard deviation. Differences between groups were assessed using a one-way ANOVA followed by a Tukey’s multiple comparisons test. This was performed using GraphPad Prism version 6.0g for Mac OS X (GraphPad Software, La Jolla California USA, www.graphpad.com). Statistical significance was considered if p≤0.05.

## Results

Oxygen saturation was successfully reduced in all hypoxic groups compared with normoxic data (**Figure 2A**).

**Figure 2:**
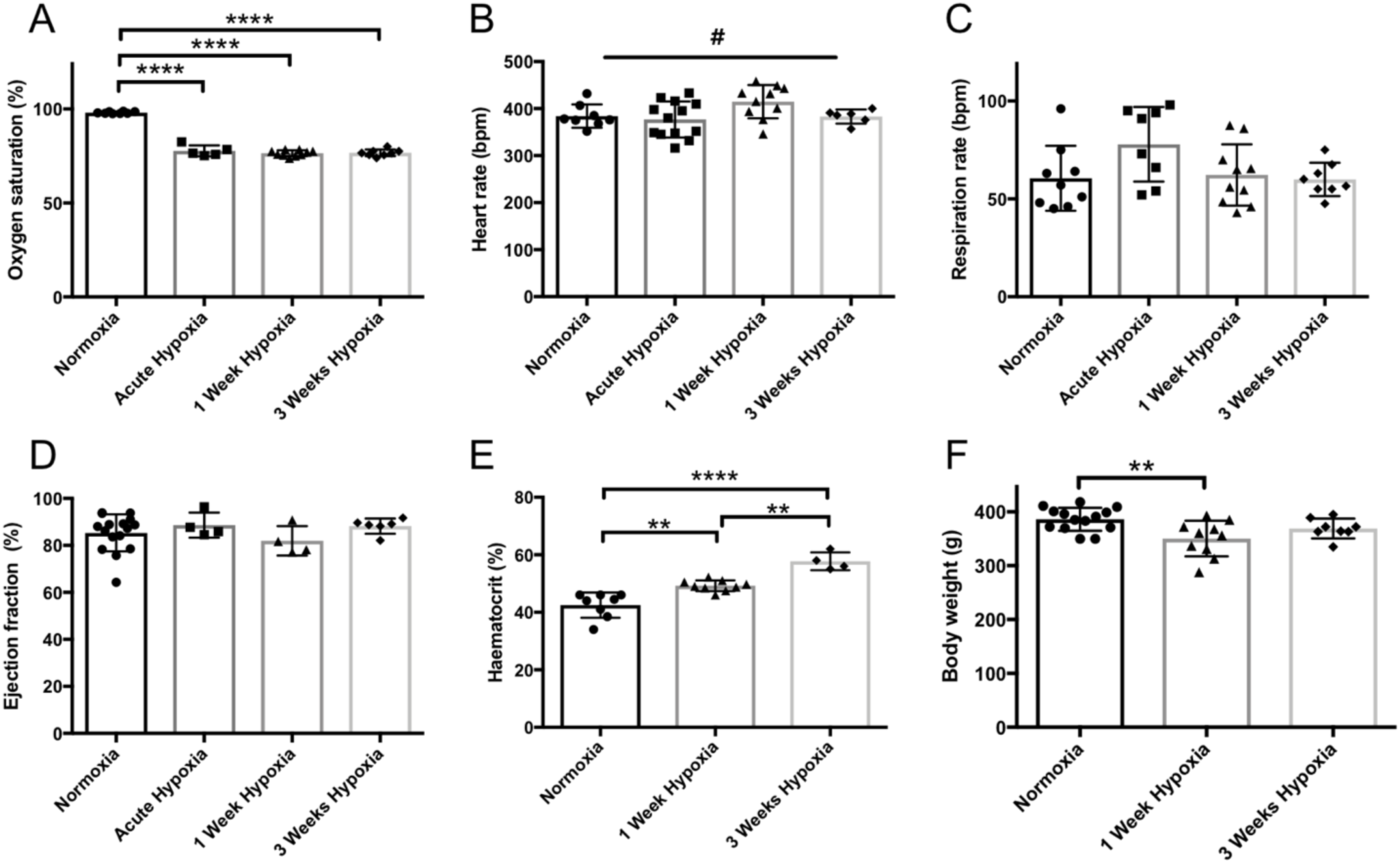
Effects of hypoxic exposure on **(A)** blood oxygen saturation, **(B)** heart rate (#p=0.051), **(C)** respiration rate, **(D)** cardiac ejection fraction, **(E)** haematocrit levels, and **(F)** body weight, all in comparison with normoxically-housed animal data; **p<0.01, ****p<0.0001.

### Physiological Effects of Hypoxia

Hypoxia did not significantly affect *in vivo* heart rate, respiration rate or left ventricular ejection fraction in any group (**Figure 2B, C, D**). However, the ANOVA for heart rate gave a p value of 0.051, and so a comparison between the 30 minute and 1-week hypoxia data should be noted (p=0.04). One week of hypoxia caused a significant increase in haematocrit compared to normoxia (49.3±0.6% and 43±2% respectively), and haematocrit in three-week hypoxic animals was significantly increased compared to one-week and normoxic values (58±2%) (**Figure 2E**); this demonstrates systemic adaptation to hypoxia over time. Animals housed in hypoxia for one week showed significantly lower body weights than normoxic animals. Following three weeks of hypoxia however, body weights were no different from controls.

### Metabolic Effects of Hypoxia

#### In vivo data

Following 30 minutes of hypoxia, animals demonstrated a significant reduction in PDH flux (50%) compared to normoxic animals (0.009±0.003 s^-1^ and 0.017±0.007 s^-1^ respectively; **Figure 3A**). In contrast, both 1 and 3 weeks of hypoxic exposure did not show significantly altered PDH flux, with values not significantly different from controls (one-week hypoxia: 0.013±0.007 s^-1^; three-weeks hypoxia: 0.017±0.011 s^-1^; normoxia: 0.017±0.007 s^-1^).

**Figure 3:**
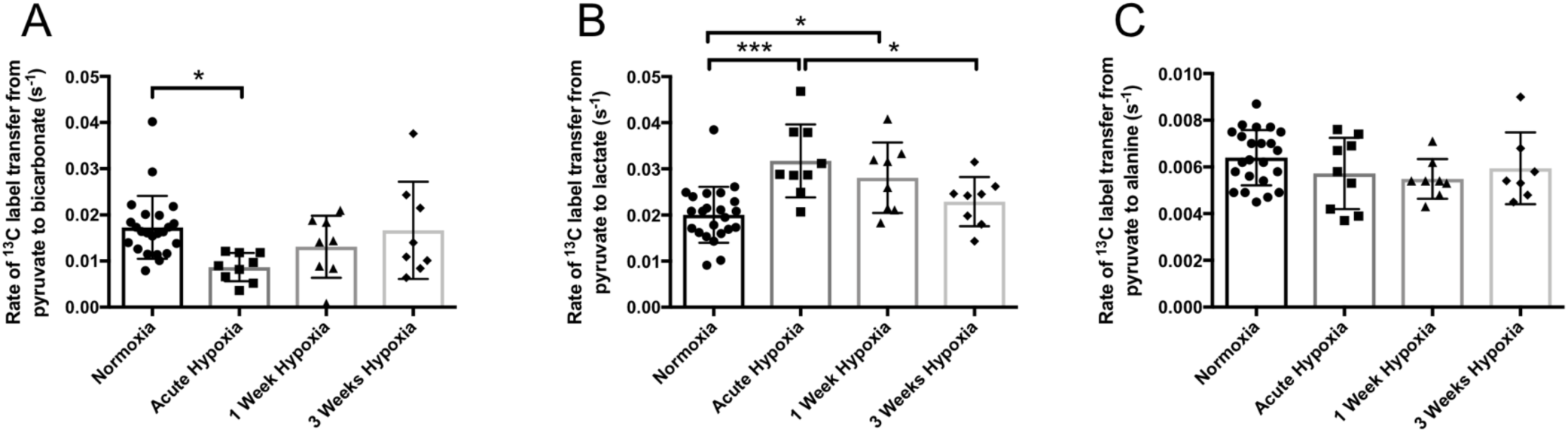
Following normoxia, 30 minutes, 1 week, and 3 weeks of hypoxic exposure, rates of HP 13C label transfer from **HP** [1-13C] pyruvate to (A) bicarbonate, (B) lactate, and (C) alanine; *p<0.05, ***p<0.001

A significant (58%) increase in HP ^13^C label transfer to lactate (**Figure 3B**), was observed in comparing 30 minutes hypoxic exposure to normoxic data (0.032±0.008 s^-1^ and 0.020±0.006 s^-1^ respectively), indicative of a short-term metabolic shift towards anaerobic metabolism. After one week of hypoxia, the unchanged PDH flux was accompanied by an increased rate of ^13^C label transfer to lactate (by 40%) compared to normoxic animals (0.028±0.008 s^-1^ and 0.020±0.006 s^-1^ respectively). No difference in flux to ^13^C lactate was observed following three weeks of hypoxia compared to normoxic data (0.023±0.002 s^-1^ and 0.020±0.001 s^-1^ respectively). No change in the rate of ^13^C label transfer to alanine was seen at any timepoint (**Figure 3C**).

### Biochemical analyses

Cardiac tissue from the one-week and three-week hypoxic groups was assessed *ex vivo*. In agreement with the unchanged PDH flux at both these timepoints, no significant differences in the protein expression levels of the regulatory cardiac PDK isoforms (1, 2 and 4) were observed (**Figure 4**). A significantly higher expression of LDH was observed in the 1-week hypoxic tissue, in line with the increased HP pyruvate to lactate conversion seen *in vivo*.

**Figure 4:**
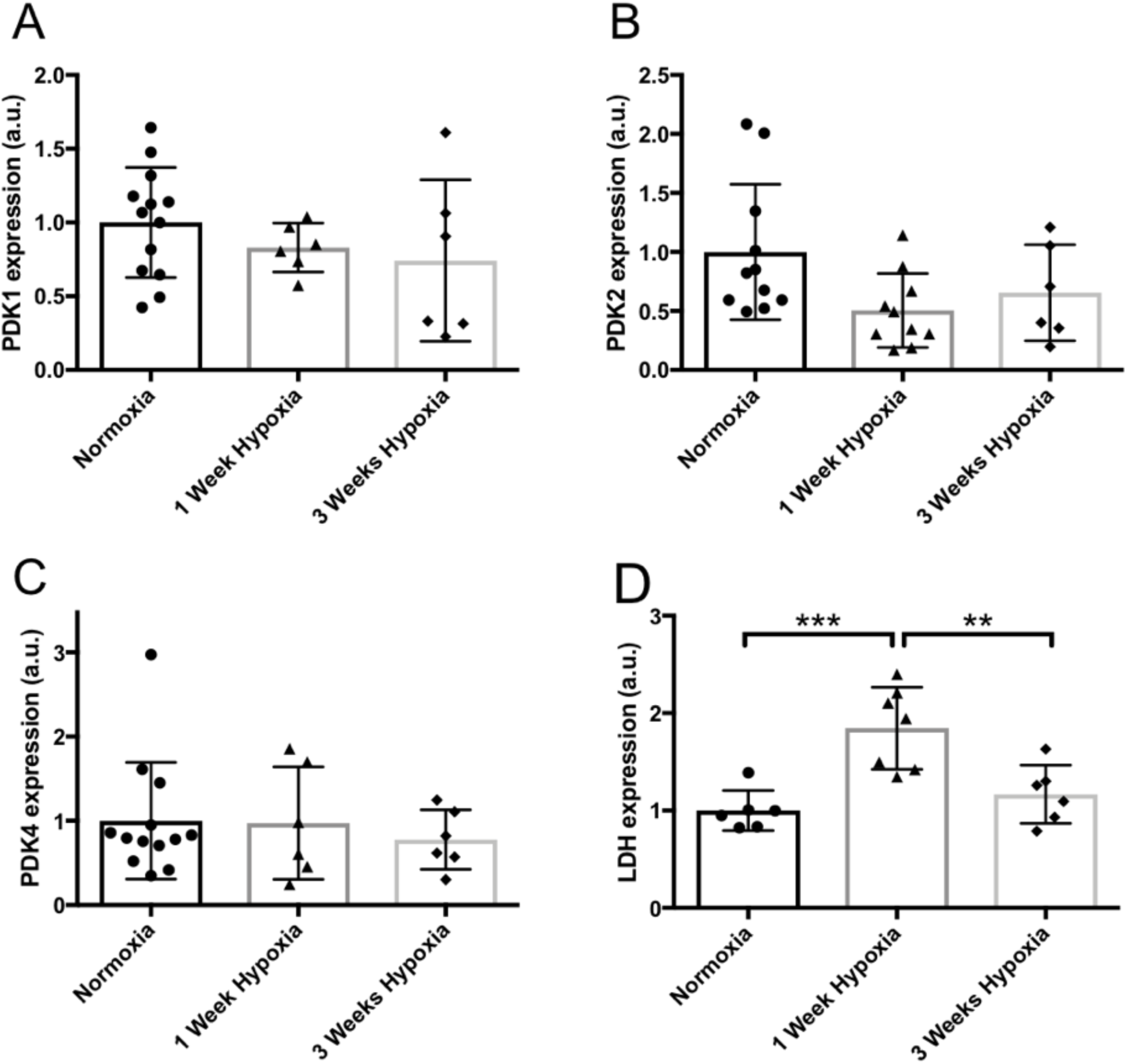
Western blot data from normoxic, 1 week hypoxic, and 3 weeks hypoxic exposure showing protein expression levels of **(A)** POK 1, **(B)** POK2, (C) POK4, and (D) LOH, **p<0.01, ***p<0.001

## Discussion

In hypoxia, metabolic changes have to occur in order for cardiac function to be maintained under these oxygen-restricted conditions. Firstly considering the response to acute hypoxia, the heart must rapidly shift metabolism towards a more anaerobic phenotype, which is characterised by increased glycolysis, increased lactate efflux^34^ and decreased oxidative mitochondrial metabolism. Indeed, in the animals exposed to 30 minutes of hypoxia, cardiac pyruvate to lactate conversion *in vivo* was significantly increased, and PDH flux significantly decreased. The rapid response that we observed, in line with the expected metabolic signature of anaerobic respiration, is likely mediated by changes in the NAD^+^/NADH ratio as a direct result of the decreased oxygen availability^35^. The reduced oxygen results in decreased mitochondrial respiration^4^, increasing NADH, inhibiting NAD-dependent dehydrogenases such as PDH and promoting NADH-dependent dehydrogenases such as LDH.

After one week of hypoxic exposure, we observed a significantly increased haematocrit level, as the animals underwent adaptation to the increasing level of hypoxia. This potentially indicates a partial adaptation to the hypoxic environment, a particularly viable suggestion when considered alongside the three-week haematocrit data, which shows an additional significant increase in haematocrit. This hypothesis of ‘interim’ adaptation is supported by a trend to increased heart rate as a compensatory mechanism to ensure sufficient systemic oxygen delivery, and a significantly reduced body weight. Similar parameters have been observed in humans adapting to altitude showing increased heart rate^36^ and a lower calorie intake^37^, the latter of has been suggested to be due to increased leptin levels^38^.

The increased haematocrit level demonstrated by our one-week and three-week hypoxic animals is a hallmark of systemic adaptation to physiological hypoxia, driven by HIF-2α-stimulated production of erythropoietin^39,40^. Glycolytic changes have been reported to be predominantly HIF-1α-regulated^41^ such as that of lactate dehydrogenase^42^, the enzyme responsible for the HP conversion we measured *in vivo*. Glycolytically derived lactate was increased in the one-week hypoxic animals, as assessed by HP pyruvate to lactate conversion, in line with significantly increased LDH expression in comparison to normoxic data. PDH flux was not decreased, which was supported by our assessment of expression levels of its PDK regulators, perhaps unexpectedly due to previous studies discussing the hypoxia-inducible nature of PDK1^43,44^. Our three-week hypoxic exposure also resulted in no metabolic differences (in the conversion of HP pyruvate to lactate or bicarbonate) in comparison with normoxic data, as supported by measures of PDK and LDH expression.

Much research has however focussed on the effect of hypoxia on PDK expression in cell culture. Kim *et al.*^43^ and Papandreou *et al*.^44^ showed upregulation of PDK1, in mouse embryonic fibroblasts following 24-72h in 0.5% hypoxia. Genetic over-activation of HIF1α increases PDK1 and 4 protein levels in muscle^45^. It has generally been assumed that this translates to the heart, in the *in vivo* setting. Equally, measured changes in these regulatory kinases have been extrapolated to mean a change in PDH activity. However, our data suggests that this may not enable comment on long-term *in vivo* cardiac hypoxia. Indeed, a study by Le Moine *et al.* demonstrated no elevation of PDK1 expression in skeletal muscle following one week of hypoxic exposure^46^. Previous studies from our group have shown that this three-week protocol of chronic hypoxia at 11% oxygen is sufficient to metabolically reprogram the heart specifically to become more oxygen efficient^5^ in ways not assessed in this study. Further, studies in animal models of hypertrophy have revealed unchanged PDH activity^47,48^ and no differences in PDK isoforms, which appeared at odds with cellular studies on hypoxia. Our data contributes to these observations and may in future help explain the situation in disease.

## Limitations

This study did not measure *ex vivo* PDH activity, which could contribute to the *in vivo* HP measures, and could be altered in spite of unchanged PDK expression. However the work by Le Moine *et al.* demonstrated that *ex vivo* skeletal PDH activity in mice exposed to one week of hypoxia was unchanged compared to normoxic animals^49^. Concomitantly, work by Atherton *et al.* demonstrated a significant correlation between *in vivo* data acquired using HP [1-^13^C] pyruvate and PDH activity assessed from *ex vivo* tissue^50^, strengthening the validity of our *in vivo* HP data. A pulse-acquire sequence was used in this study, and data acquired using a surface coil. Future work could involve implementing a more elegant acquisition protocol^51^ to provide more information on regional hypoxia within the heart.

Finally, normoxic animals were imaged using 100% oxygen, which, although common procedure in preclinical animal studies, may exacerbate the differences we have seen here. Future studies could include anaesthesia at a lower oxygen percentage.

## Conclusion

In conclusion, we have demonstrated the ability of HP [1-^13^C] pyruvate to non-invasively assess metabolic changes in the healthy heart in response to three lengths of exposure to hypoxia. This could therefore be a viable technique for assessing hypoxia in a wide range of diseases and in response to therapy.

## Acknowledgements

The authors would like to thank Dr. Louise Upton and Prof. Mary Sugden for the kind provision of the pulse oximeter and a primary antibody for PDK4 respectively. L.L.P would also like to thank Richard and Jocelyn Le Page for technical assistance in preparing the manuscript, and Asst. Prof. Myriam Chaumeil for valuable discussions.

## Abbreviations

HP: hyperpolarized
PDH: pyruvate dehydrogenase
LDH: lactate dehydrogenase
BOLD: blood-oxygen-level dependent
PET: positron emission tomography
PDK: pyruvate dehydrogenase kinase
HIF: hypoxia-inducible factor

